# scIntegral: A scalable and accurate cell-type identification method for scRNA-seq data with application to integration of multiple donors

**DOI:** 10.1101/2020.09.17.301911

**Authors:** Hanbin Lee, Chanwoo Kim, Juhee Jeong, Keehoon Jung, Buhm Han

**Author notes:** **Emails**, Hanbin Lee, Chanwoo Kim, Juhee Jung, Keehoon Jung, Buhm Han. **Corresponding Author**, Buhm Han, Department of Medicine, Seoul National University College of Medicine, 103 Daehak-ro, Jongno-gu, Seoul 03080, South Korea. These authors contributed equally (Hanbin Lee, Chanwoo Kim).

## Abstract

We present *scIntegral*, a scalable and accurate method to identify cell types in scRNA data. Our method probabilistically identifies cell-types of the cells in a semi-supervised manner using marker list information as prior. scIntegral is more accurate than existing state-of-the-art methods, reducing the error rate by up to three-folds in real data. scIntegral can precisely identify very rare (<0.5%) cell populations, suggesting utilities for *in-silico* cell extraction. A notable application of scIntegral is to systematically integrate scRNA-seq data of multiple donors with strong heterogeneity and batch effects. scIntegral is extremely efficient and takes only an hour to integrate ten thousand donor data, while fully accounting for heterogeneity with covariates. Many previous methods focused on integrating multi-sample data in the cluster level, but it was challenging to quantitatively measure the benefit of integration. We show that integrating multiple donors can significantly reduce the error rate in cell-type identification, when measured with respect to the gold standard cell labels. scIntegral is freely available at https://github.com/hanbin973/scIntegral.

## Introduction

Single-cell RNA sequencing technology (scRNA-seq) enabled dissection of expression profile at an individual cell level. A typical analysis procedure for scRNA-seq data begins with the use of an unsupervised clustering algorithm such as tSNE to project data in 2D. Based on this projection, the cell type of each cluster is manually determined using the expressions of known marker genes. However, this commonly used pipeline is prone to errors and extremely time-consuming. Unsupervised clustering algorithms are optimized to visualize the high-level structure of the data, but do not consider the marker information during clustering. For this reason, the resulting 2D projection often gives unclear boundaries between the clusters, which makes manual curation difficult. In addition, marker signatures frequently correspond to more than one cell-type introducing ambiguities in manual curation.

To address this challenge, recent studies have proposed probabilistic semi-supervised approaches (1,2). These algorithms directly utilize the knowledge of marker genes to probabilistically assign cells to cell-types, rather than using markers only to label the clusters obtained through unsupervised clustering. SCINA uses a Gaussian Expectation-Maximization (EM) algorithm after normalization of the expressions (1). CellAssign similarly uses an EM algorithm but directly models the expression counts with the Negative Binomial distribution (2). These methods can provide useful information to complement the common analysis pipeline based on 2D projection; with these methods, one can disclose the identity of each single cell, determine the label of a group of cells defined in the 2D projection, assess proportions of celltypes, and perform differential gene expression (DEG) analysis based on cellular identities. However, these methods still have limited accuracy.

Here we present *scIntegral,* a method that accurately assigns cells to cell-types in scRNA data. scIntegral has superior accuracy compared to previous methods, while obtaining high robustness, model generality, and speed. Similar to the existing method (2), our method begins with a general model that can incorporate covariate effects and overlapping markers between types. To maximize accuracy, however, scIntegral uses two strategies. First, scIntegral constructs an informative prior to utilize marker information more effectively. Informative prior enables smart initialization so that the optimization algorithm can avoid local optima and properly reach the global optimum. Second, scIntegral is equipped with a custom gradient computing routine. By avoiding unnecessary matrix multiplication and storage, gradient calculation is faster up to 3folds and has a much less memory requirement compared to automatic differentiation schemes provided by deep learning libraries.

Our method is high-resolution and accurate. In the real benchmarking datasets, scIntegral outperformed existing methods in accuracy. For example, in the stem cell dataset of Koh et al. (3), scIntegral achieved 96.8% median accuracy in classifying 446 cells into 8 cell-types, while SCINA and CellAssign only achieved 80% and 89.9% median accuracy, respectively. Thus, the error rate was reduced by three-folds. The variance of accuracy was also dramatically reduced compared to the competing method, as all independent runs of our method reached the global optimum. Our method particularly excels in identifying rare populations. If a cell population is very rare (e.g. <0.5%), typical unsupervised clustering algorithms have difficulties in identifying it as a separate group. The manual curation via human eyes is also not optimal for finding a small population. When applied to the pancreas datasets(4,5) (4,679 cells), our method identified Schwann cells correctly that consisted of only 6 cells (~0.1% of the data). This high-resolution accuracy suggests that our method can be used for *in-silico* extraction of target cells that exist in a very small amount in the data.

A notable application of scIntegral is to systematically integrate scRNA-seq data of multiple donors. Each individual donor sample presents a snapshot of the cell ecosystem that may change temporally. Thus, like many biological experiments, merging multiple donor samples together can suppress the individual-specific variations in the data and reveal shared components. Although the benefits of integration are apparent, combining multiple donor samples is extremely challenging. A simple merging of cell data would not work because of the heterogeneity between donor samples. Donor samples may present heterogeneity caused by different biological states as well as heterogeneity caused by different experimental conditions and technological platforms, if the samples come from multiple studies.

There have been active efforts to develop methods to integrate scRNA data of multiple donor samples. Many methods focused on integrating data in the cluster level. Seurat chooses one sample as a reference and aligns the clusters of the other samples to the reference(6). scAlign uses a deep neural network to project multiple sample data to a shared space and performs clustering in that space(7). Harmony integrates multiple datasets in the principal component spaces by calculating the most likely alignment(8). The limitation of these methods is that although they could integrate multiple donor samples, it was difficult to prove how good the final alignment was, because the quality of clustering can hardly be quantified. Another category of methods is normalization(9,10). These methods try to regress out the effect of between-sample heterogeneity from the cell count data. The normalized data can then be integrated together, as if all cells came from a single donor. However, it has not been shown if the normalization could effectively remove all heterogeneity between donors.

Because of its general model and extreme efficiency, scIntegral can easily integrate scRNA data of multiple donor samples during its probabilistic cell-type assignment procedure. Because we built our method with scalability in mind, in theory, our method can integrate and analyze ten thousand donor samples together in less than an hour. Since the quality of cell-type identification is directly quantifiable by comparing to the experimentally curated cell labels, we were able to systematically measure the benefit of integration. We obtained the liver scRNA-seq dataset that consists of 8,444 cells from 5 donors. When we analyzed each donor separately, the overall celltype identification accuracy was 88.7%. However, when we integrated all donors from the two studies together using our method, the accuracy increased to 93.1%. Thus, the error rate was reduced by 40%. We also obtained two human pancreas datasets(4,5), one consisting of 2,394 cells from 10 donors and another consisting of 2,285 cells from 4 donors. In this dataset, our method already had a very high accuracy (97.3%) even when analyzing single donors, but the error rate was reduced from 2.7% to 2.2% by integrating all 14 donors. Overall, these demonstrated that the donor data supported each other, and the integration of all donors was the most effective. That is, like other biological experiments, scRNA analysis could benefit from integration of multiple donors and experiments. We note that the accuracy dropped to 60.0% when we simply merged all cell data without covariates, which emphasizes that a proper integration is important.

There were many cell-type identification methods, but our method differs from them. SCINA(1) and CellAssign(2) are similar semi-supervised methods to ours. SCINA cannot incorporate covariates, and therefore cannot be applied to integrate multiple donors. CellAssign can incorporate covariates, but integration of hundreds of samples is computationally infeasible. We show that our method is more accurate and robust than SCINA and CellAssign in the analysis of real data. Another category of methods are supervised methods that use not only the marker list information but also the expression of the markers that are present in existing databases. Although supervised methods can utilize more from prior knowledge, they have limitations that the methods cannot identify cell types whose expressions are not in the databases. Indeed, for human embryogenic stem cell dataset(3) used in our analysis, there was no appropriate and complete database that contained expressions of all candidate cell types. Moreover, supervised methods cannot take advantage of data integration of multiple donors, because they process each donor data separately by comparing it to the database.

Our method is publicly available at https://github.com/hanbin973/scIntegral.

## Materials and Methods

### The scIntegral model

#### Linear Negative binomial model

Negative binomial regression is a famous regression method for overdispersed count data. The following model is fitted.

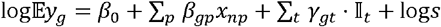

Here, *β*_0_ is a constant intercept, *β_gp_* are covariate coefficients, *γ_gt_* are cell-type specific expression coefficients and s is a size-factor. *x_np_* and 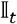 are covariate and cell-type indicators respectively. Covariates include donors and technological platforms.

Under this model, we can express the mean expression count *μ_ngt_* of gene *g* in the cell *n* of celltype *t* as

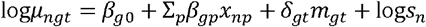

Let *y_ng_* be the observed expression count of gene *g* in cell *n*. We assume that *y_ng_* follows a negative-binomial distribution. Here, *β*_0_ is the constant, *β_gp_* are covariate coefficients, *δ_gt_* are celltype specific expression coefficient and *s_n_* is the size-factor. *m_gt_* is 1 if gene *g* is a marker of celltype *t* and otherwise 0.

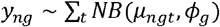

Existing theoretical literature predict that the dispersion of the expression count is determined by the gene regulation mechanism(11,12). By assuming that gene regulation mechanism is independent of the celltype for the same gene, we model the dispersion to be dependent only on the gene *g* but not celltype *t*, thus *ϕ_g_*.

The full likelihood is

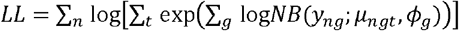

#### Initialization through an informative prior based on marker specification

The log-likelihood function of a mixture model generally does not have a unique optimum. Multiple local optima can degrade the stability when fitting the model. For example, we found that existing method requires multiple runs to reach a proper solution to the likelihood optimization problem. If additional information about an optimum is provided, such information can be used to improve the convergence of the model.

In our model, we designed an initialization scheme that maximally deploys the input marker information. When a gene is a marker of a given cell-type, *δ_gt_* is initialized so that it is strictly larger than a threshold that is larger than 0. We found that setting this threshold to 2 generalizes over different datasets well. Additionally, we modified the spike-and-slab prior widely used in sparse optimization literatures as follows:

- 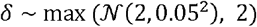

We then initialized the parameter *β* as follows:

- 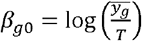
- *β_gp_* = 0 for *p* = 1,...,*P*

where 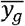 is the mean of the row *g* of the expression matrix *Y*.

#### Smart calculation of gradient

The goal of scIntegral is to probabilistically assign cells to cell-types. Conversely, this can be done by estimating the correct parameters of the model, because once the parameters (*γ_gt_,ϕ_g_,β_gp_*} are known, it is straightforward to calculate the posterior probability of the cell-types by Bayes rule.

Unfortunately, there is no closed-form solution to finding parameters that maximizes the likelihood. Therefore, numerical optimization is required. To this end, we use a deep learning framework in PyTorch. Deep learning frameworks provide a package of optimization methods that work by gradient calculation and back-propagation, which can be conveniently used for any model even if the model is not a deep neural network. Indeed, a previous study (2) also used a deep learning framework for celltype identification.

In a deep learning framework, the gradient is calculated by automatic differentiation. Automatic differentiation is a fast and a general gradient computing algorithm that is implemented in famous deep learning libraries. PyTorch, the backend of our scIntegral implementation, also supports this feature as Autograd. Thus, the log-likelihood of scIntegral can be differentiated using Autograd. However, Autograd ignores the specific structure of the optimization target because it computes the gradient using the famous chain-rule of calculus.

To increase speed and scalability of our method, we analytically derived the gradient for each of the unknown parameters by carefully examining the log-likelihood of scIntegral. The gradient of the loglikelihood respect to the cell type specific parameter *δ_gt_* is

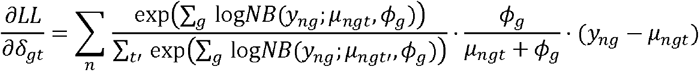

The gradient of the log-likelihood respect to the dispersion parameter *ϕ_g_* is

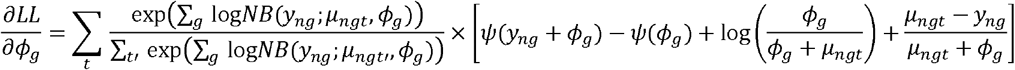

The gradient of the log-likelihood respect to the covariate effect *β_gp_* is

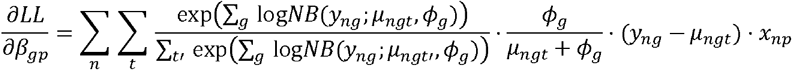

The remaining details about the derivation of the gradients can be found in the **Supplemental_Notes.pdf**.

After deriving the closed form formula for the gradient, we implemented a routine that demands a minimal number of matrix multiplications and does not rely on the chain rule (**Figure 3a**). This implementation has two advantages over using Autograd. First, there are fewer matrix multiplications compared to Autograd. Autograd computes the gradient of the optimization target layer by layer and multiplies all the gradient of each layer in a fixed order. This can be unnecessarily demanding because the total number of operations depends on the multiplication order of the matrices. Second, our implementation demands less memory storage compared to Autograd. Storing large matrices is not only memory consuming but also time consuming. Autograd stores intermediate values during target optimization to avoid computing the same quantity multiple times. This, however, comes at the cost of large memory overhead. Our implementation minimized the number of intermediate matrices required to avoid unnecessary memory storage. Overall, this technical advance in gradient calculation has given our method extremely high efficiency and scalability.

#### Implementation

scIntegral is coded in Python 3.7 using PyTorch v1.4. The required packages are numpy, scipy and pandas which can be straightforwardly installed. The method uses GPU, if installed, and CPU otherwise. The package is available at the github repository.

### Accuracy measurement

True cell-type labels were provided in the metadata of each dataset. We then measured the accuracy of each method by dividing the number of cells in which the true cell-type label and the assigned label by the method matched with the total number of cells in each dataset.

### Comparison with existing methods

We compared our method to SCINA(1) and CellAssign(2). For SCINA, we installed and used the current version (v1.2.0) on CRAN. For CellAssign, we installed and used the current version (v0.99.16) coded in R-TensorFlow (v2.2.0) with default options. CellAssign uses the EM algorithm where M step is optimized using the Adam optimizer with Autograd. The current implementation of SCINA and CellAssign does not run on GPU (as of 06/20/2020), but future versions may.

In our comparison, we used the same marker sets, covariates, and cell size factors for all methods. For liver dataset, we computed cell size factors using computeSumFactors(13) function from the scran R package. For other datasets, we used pre-calculated cell size factors included in the DuoClustering2018 R package. We used the same stop criterion (<0.01% change in LL) for CellAssign and scIntegral. For SCINA, the default convergence criterion was used. All the analysis used the same computer resource (1 CPU = 4 threads). For all methods, we included the unknown category to capture unknown cell-types.

### Benchmarking hardware

All benchmarking took place on an Intel^®^ Xeon^®^ Gold 6136 CPU (3GHz). We used a single CPU and limited the number of threads to four using the taskset command in CentOS 7 operating system. A single Nvidia^®^ RTX^®^ 2080 was used in GPU mode.

### t-SNE

In all our datasets except for human pancreas(4,5), meta information of the samples including the t-SNE coordinates were available. To maintain comparability with the previous methods, we directly adopted the t-SNE coordinates from these metadata to plot **Figure 3a-l**. For human pancreas, t-SNE coordinates were obtained using Seurat(14).

### Number of donors analyzed per hour

This quantity was calculated as follows. The hour in seconds (3,600 seconds per hour) was divided by the runtime for each dataset. Finally, the number of donors was multiplied.

#### Embryonic stem cell scRNA data

The Koh et al. dataset consists of 446 human embryonic stem cells (hESCs) at various stages of differentiation. We extracted the data from the R package DuoClustering2018, which can be installed using Bioconductor package manager. The dataset contains 9 cell types. Among them, we used 8 cell types with both scRNA-seq data and bulk RNA-seq data, which are hESC (day 0), Anterior Primitive Streak (day 1), Mid Primitive Streak (day 1), DLL1+ Paraxial Mesoderm (day 2), Lateral Mesoderm (day 2), Early Somite (day 3), Sclerotome (day 6), Central Dermomyotome (day 5). Koh et al. annotated the cell types through fluorescence activated cell sorting (FACS). We defined marker genes for each cell type using the bulk RNA-seq data following the same procedure described in Zhang et al. 2. Briefly, for each gene, we sorted the N=8 types in ascending order based on the mean expression level. We then calculated log fold change between two consecutive types in this order. We then chose the maximum value among the N-1 log fold change values. After calculating this maximum value for all genes, we used genes with the maximum value in the top 20th percentile as marker genes.

#### Human liver scRNA data

The MacParland et al. liver dataset consists of 11 cell types with 8,444 cells collected from 5 patients. We extracted the data from the R package HumanLiver, which can be downloaded from https://github.com/BaderLab/HumanLiver. After clustering cells, MacParland et al. determined the identity of each cluster using known gene expression profiles. We mapped 20 discrete cell populations identified by them to 11 unique cell types for our analysis (Hepatocytes, ab T cells, Macrophages, Plasma cells, NK cells, gd T cells, LSECs, Mature B cells, Cholangiocytes, Erythroid cells, Hepatic Stellate Cells). The mapping was obtained using ClusterNames() function. We applied scIntegral and CellAssign to this dataset using the marker genes described in the supplementary materials of Zhang et al.. We included patient information as covariates in both methods.

#### Human pancreas dataset

Two datasets with total 14 donors were from Muraro et al. and Segerstolpe et al. (4,5). Total 4,679 cells were included. The data were openly available at GEO85241 (https://www.ncbi.nlm.nih.gov/geo/query/acc.cgi?acc=GSE85241) and E-MTAB-5060 (https://www.ebi.ac.uk/arrayexpress/experiments/E-MTAB-5060/). The markers were retrieved from the PanglaoDB(15). Muraro et al. used the CEL-seq2 platform and Segerstolpe et al. used the SMART-seq2 platform. Each dataset consists of 2,285 and 2,384 cells from 4 and 10 donors respectively. The original data contained 14 cell populations (alpha, gamma, acinar, ductal, beta, delta, activated stellate, inactivated stellate, macrophage, endothelial, Schwann, mast and epsilon). Because the PanglaoDB did not have separated markers for activated and inactivated stellate cells, we merged the two populations into a single population.

#### PBMC 4k scRNA data

We obtained PBMC 4k (peripheral blood mononuclear cell) dataset, namely Zhengmix4eq, from the R package DuoClustering2018. This dataset is a mixture of 3,994 FACS purified PBMC cells of 4 cell types, which are B-cells, CD14 monocytes, naive cytotoxic T-cells and regulatory T-cells. We used only the genes of which mean expression (log-normalized count) value across all cells was in top 30th percentile. We defined marker genes for each cell type in the procedure similar to Koh et al. as follows. We sorted N=4 types in ascending order based on the mean expression, and calculated log fold change between consecutive types in this order. Among the N-1 log fold change values, we chose the maximum. We then used the genes of which the maximum value is in the top 30th percentile as marker genes.

#### PBMC 68k scRNA data

We obtained PBMC 68k (peripheral blood mononuclear cell) dataset from the 10x genomics data download page (https://support.10xgenomics.com/single-cell-gene-expression/datasets). This data consists of total 68,579 cells. Unlike other datasets described above, the cell labels of this data do not exist. In the original study(16), the cell labels were determined computationally using a FACS purified PBMC sample. Therefore, we only used this dataset to obtain computation time and did not measure the classification accuracy. Markers were retrieved similarly to the PBMC 4k dataset. 10x genomics provides a mean expression level of purified PBMC cells. We sorted N=11 types in ascending order based on the mean expression, and calculated log fold change between consecutive types in this order. Among the N-1 log fold change values, we chose the maximum. We then used the genes of which the maximum value is in the top 0.2th percentile as marker genes (total 127 markers).

## Results

### Linear negative binomial models can explain mRNA expression variability

scIntegral takes the raw expression counts as inputs to predict the cell type of each cell. Both theoretical and empirical research have shown that expression counts follow a Poisson or a negative binomial distribution(12,17,18). Furthermore, recent researches showed that the expression count of a given gene depends on its regulatory mechanism(11,12). Inspired by these observations, scIntegral models the expression counts of a heterogeneous cell population using a linear negative binomial mixture model (**Materials and Methods**). Negative binomial distribution is a generalization of the Poisson distribution that can account for over-dispersion which is frequently found in immune related genes(18). Each negative binomial component is then modeled by a linear combination of parameters. One might argue that scRNA-seq data is too complex to be modelled with a simple linear modelling strategy. Here we show that linear combination of size factor, cell type indicators and donor indicators together can explain a large portion of the variability observed in the expression count data (**Materials and Methods**). When the linear model was applied to human liver and human pancreas datasets, the median explained variance (pseudo-R^2^) among marker genes were larger than 60% in both datasets showing that a linear model is an effective modelling strategy for scRNA-seq data (**Figure 1a and 1c**). Based on this model, scIntegral infers the parameters of each mixture component and computes the posterior probability of the cell type for each cell. Note that existing differentially expressed gene analysis (DEG) methods such as DESeq2(19) and edgeR(20) also adopt linear negative binomial models.

**Figure 1.**
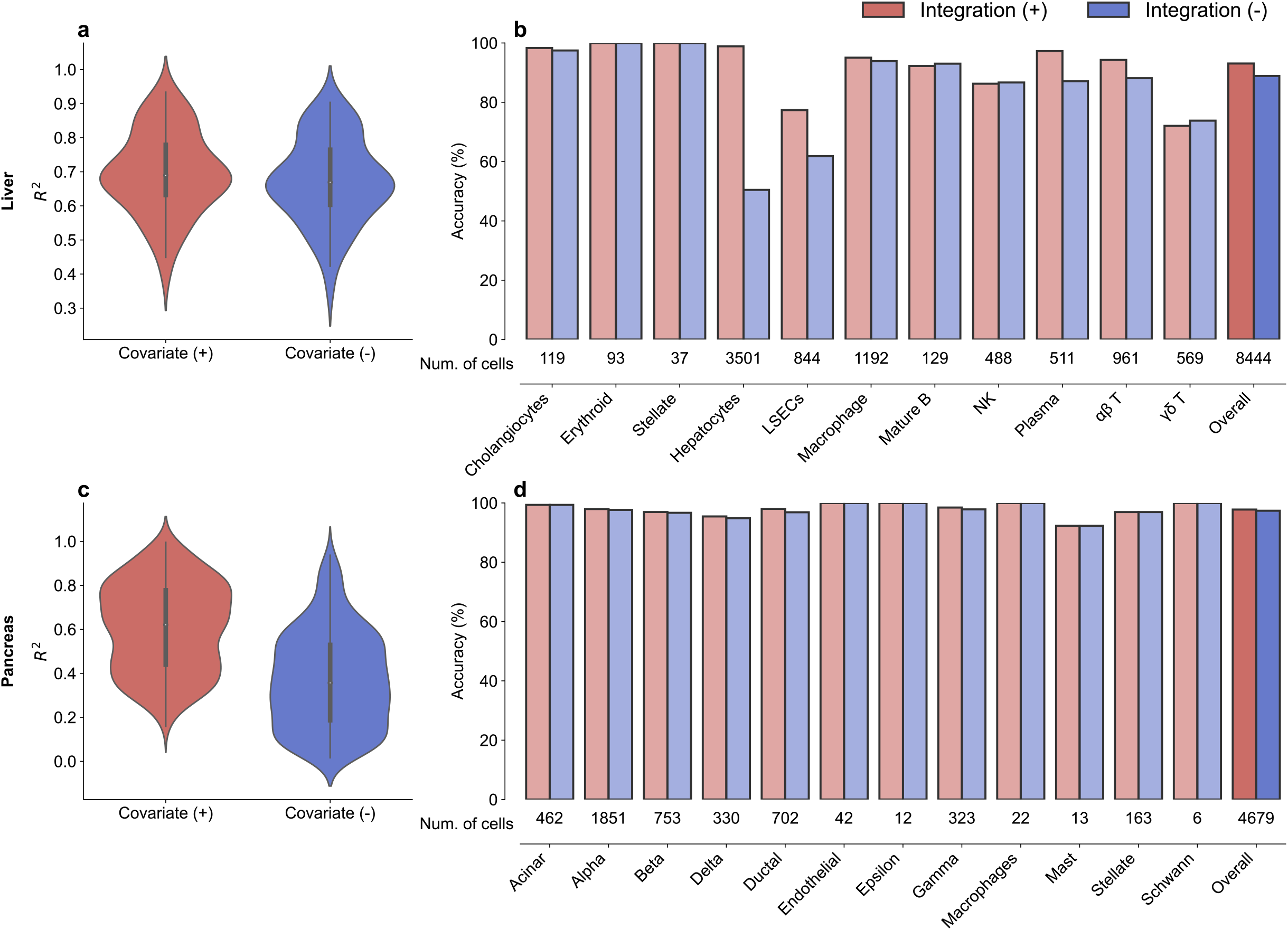
Model fitness of linear models and accuracy improvement after integration. **a.** Pseudo R^2^ of negative binomial models with and without covariates in the human liver dataset(21). **b.** Classification accuracy of each cell type using scIntegral with and without integration in the human liver dataset. **c.** Pseudo R^2^ of negative binomial models with and without covariates in the human pancreas datasets(4,5). **d.** Classification accuracy of each cell type using scIntegral with and without integration in the human pancreas datasets.

### scIntegral properly handles confounding effects due to technical variations

Although large datasets containing multiple experiments and donors can lead to more precise prediction of the cell types and estimation of model parameters, heterogeneity underlying the data can confound the analysis. We first measured the variance explained by the information of the cell donor. We applied negative binomial regression to the human liver(21) and human pancreas datasets(4,5) (**Materials and Methods**). These data consist of five and fourteen different donors respectively. The negative binomial regression was fitted with and without donor indicator variables. We see that adding the donor indicators increased the model R^2^ up to 20% (**Figure 1a** and **1c)**. The p-values of the indicator variables were also significant after correcting for multiple testing (**Supplemental_Table_S1.xlsx and Supplemental_Table_S2.xlsx**). Therefore, we conclude that when integrating multiple experiments, technical variation must be explicitly accounted. In the liver and pancreas datasets, scIntegral demonstrated high prediction accuracy of 93.0% and 97.8% respectively, after appropriately correcting for the donor effects (**Figure 1b and 1d**). We visualized the result through a t-SNE plot by comparing scIntegral assigned results to true cell type labels. In both liver and pancreas datasets, misclassified cells were hardly visible (**Figure 2**).

**Figure 2.**
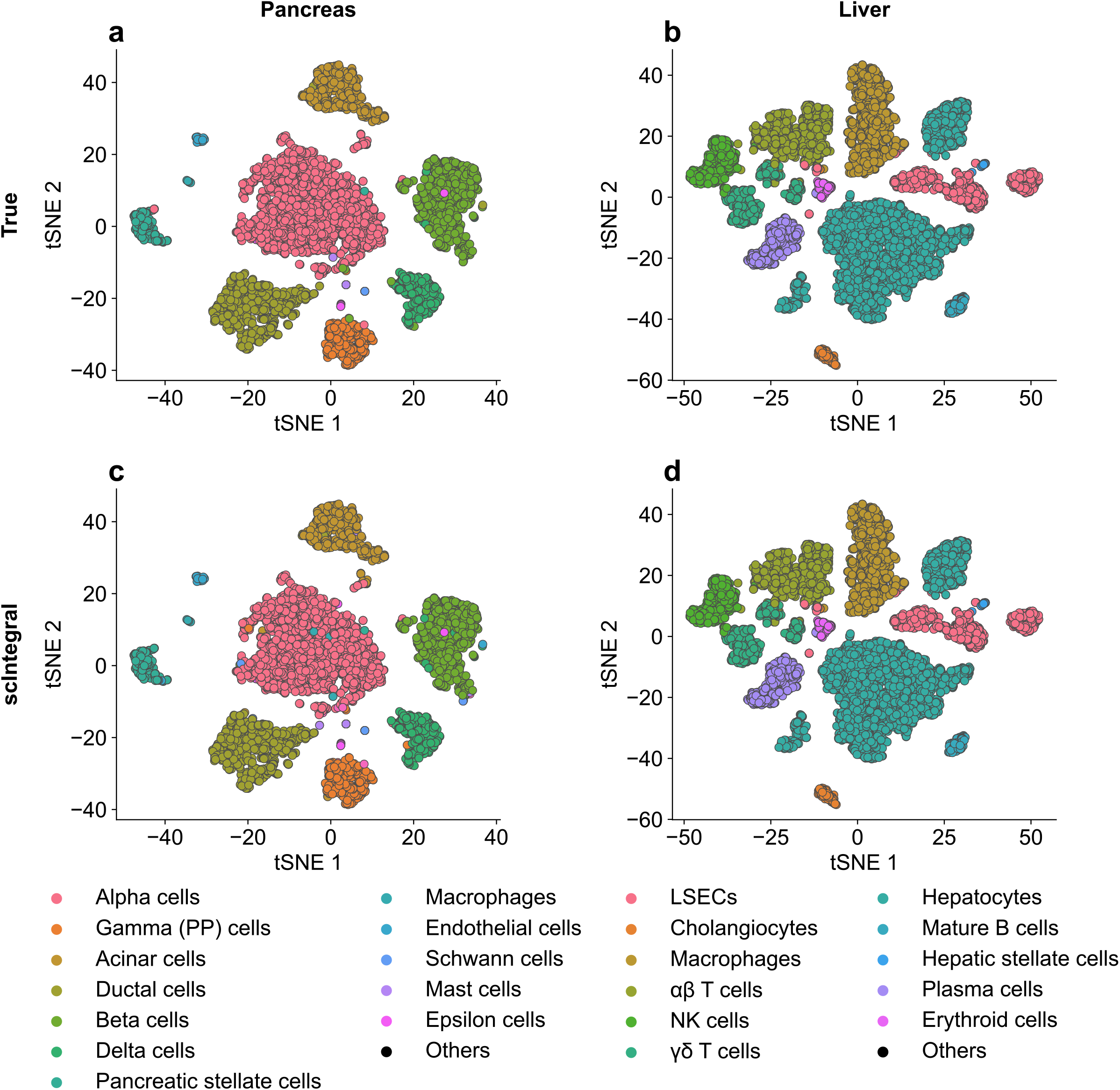
t-SNE plots of true labels and scIntegral assignments. **a.** t-SNE plot of the human pancreas datasets labeled by the true cell type labels. **b.** t-SNE plot of the human liver datasets labels by assignment result of scIntegral. **c.** t-SNE plot of the human liver dataset labeled by the true cell type labels. **d.** t-SNE plot of the human liver dataset labeled by assignment result of scIntegral.

### Integrating multiple experiments improves classification accuracy

scIntegral can efficiently integrate multiple datasets from different technological platforms, fully leveraging the opportunities of large-scale data. It is well known that in many statistical models, larger datasets can help obtain more precise estimates of the parameters and lead to improved outcome. Similarly, in many biological experiments, it is common to combine multiple samples or replicates to suppress individual-specific variations in the analysis. However, in scRNA analysis, it was unclear what the benefit would be from integrating multiple donor samples. Some studies tried integrating multiple data in the cluster level, and used specific metric to measure the existence of remaining batch effects(8). However, the benefit of multi-sample integration has not been quantified using experimentally validated data.

Here, we wanted to objectively quantify the benefit of multi-sample integration in cell type identification. Since the experimentally curated cell labels such as from FACS sorting can provide the gold standard validation, we have an opportunity to quantify the effect of integration in terms of the identification accuracy. Specifically, we show that analyzing multiple experiments together has advantages over analyzing each donor and aggregating the results afterwards. We divided the liver data into five sub-data based on its donor label. Next, we applied scIntegral to each subdata to infer cell-type labels. Compared to the cell-type assignment applied to the whole data, the results applied to each sub-data from individual donors were much less accurate (**Figure 1b and 1d**). The overall accuracy when analyzed separately was 88.7% compared to 93.1% in the combined analysis. Thus, the error rate was reduced from 11.3% to 6.9% by integrating data of 5 donors. This analysis was repeated in the pancreas dataset with fourteen different donors from two different technological platforms, SMART-Seq2 and CEL-Seq2. In this dataset, the accuracy of our method in the donor-separated analysis was already as high as 97.3%. In the combined analysis integrating data of 14 donors, the overall accuracy increased to 97.8% reducing the error rate from 2.7% to 2.2%. Overall, these showed that integrating multiple donors could decrease the cell-type classification error rate. We can expect that, as more donor data can be integrated, the cell-type identification accuracy can be further improved.

### scIntegral can identify rare cell types

scIntegral can identify very rare cell populations that occupy less than a single percentage in the whole data. In the liver dataset, hepatic stellate cells, erythroid cells, cholangiocytes and mature B cells each occupy less than 5% of total population (**Figure 1b, 1d and Supplemental_Table_S3.xlsx**). Surprisingly, scIntegral classified these rare cells with more than 95% accuracy on average (**Figure 1b, 1d and Supplemental_Table_S3.xlsx**). In the pancreas dataset, more extreme cases were found. Cell-types such as epsilon cells and Schwann cells each occupied less than 0.5% of the whole cell population (**Figure 1b, 1d and Supplemental_Table_S4.xlsx**). scIntegral successfully identified these rare cell types with 100% accuracy (**Figure 1b, 1d and Supplemental_Table_S4.xlsx**). For example, scIntegral was able to exactly specify the 6 Schwann cells. This high accuracy for rare populations suggests that our method can be used for the in-silico extraction of very rare cells from the whole sample.

### Handling large number of samples and donors

To fully leverage the advantages of data integration, scIntegral was designed with scalability in mind. Instead of using the autograd function of deep-learning frameworks, we designed a fast optimized gradient computation routine (**Figure 3a, Methods**). Using highly optimized matrix operations, scIntegral achieves significant reduction in computation time.

**Figure 3.**
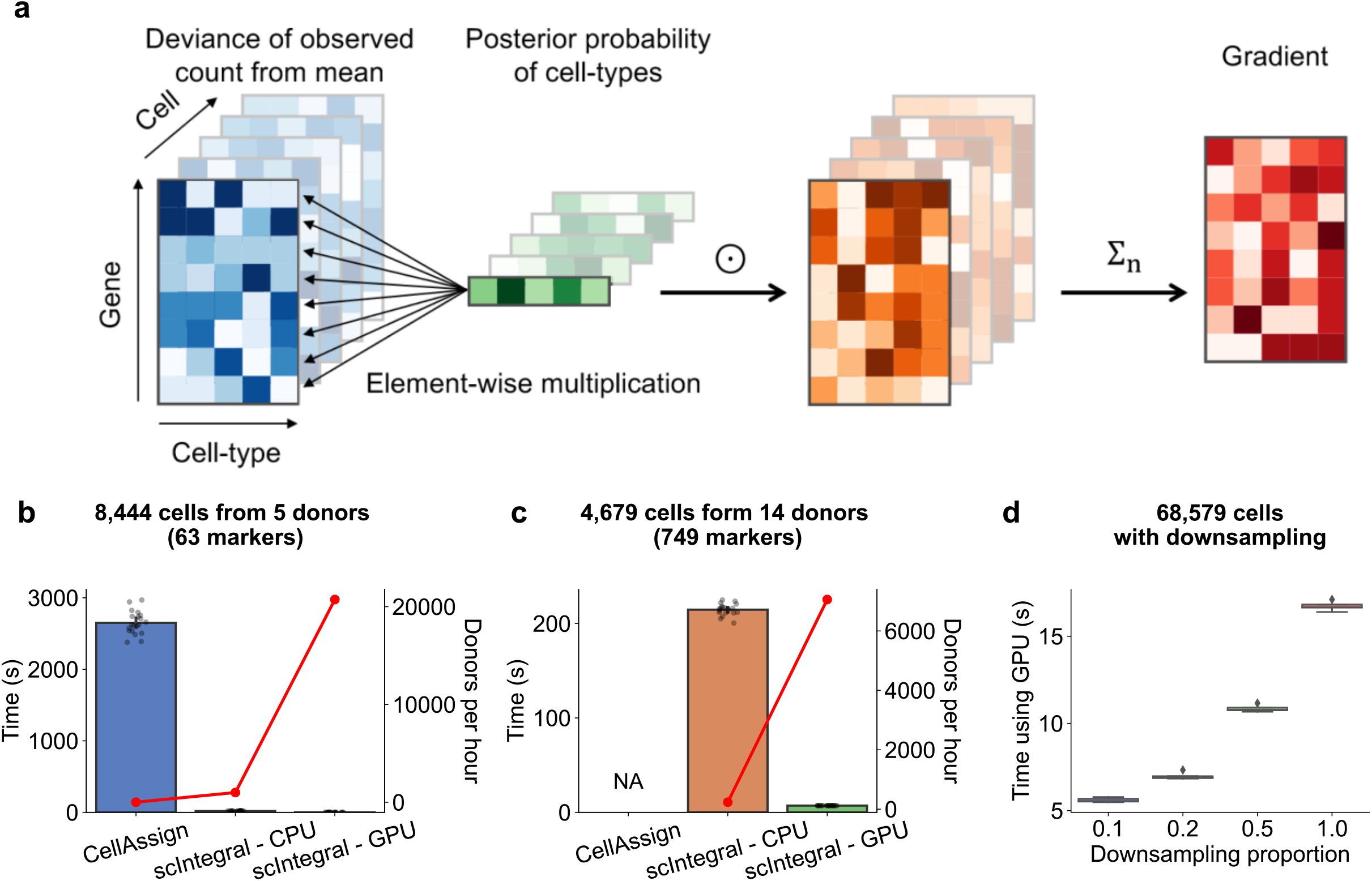
Speed benchmark of scIntegral. **a.** A graphical illustration of scIntegral’s optimized gradient computing routine. All the computations are based on highly optimized matrix routines provided by linear algebra libraries. **b.** Computation time and number of donors processed per hour of CellAssign(2) and scIntegral in the human liver dataset. **c.** Computation time and number of donors processed per hour of CellAssign(2) and scIntegral in the human pancreas datasets. **d.** Computation time of scIntegral in the PBMC 68k(16) dataset with different down-sampling proportions.

We measure the computational efficiency of scIntegral in human liver and pancreas datasets. In the liver dataset, CellAssign took 2625.0 seconds while scIntegral took only 20.0 seconds in 4-core CPU which was a 131.25 times reduction in runtime (**Figure 3b**). This is translated into processing 900 donors in an hour. In the pancreas dataset, CellAssign failed to run in a computer with 16GB memory possibly due to large number of markers (**Figure 3c**). However, scIntegral ran successfully taking 216 seconds, equivalent to processing 234 donors per hour. scIntegral supports GPU computation that can further increase the scalability of the algorithm. Using a single GPU, scIntegral took only 0.9 and 6.9 seconds in the liver and the pancreas datasets respectively. This is equivalent to processing 21,000 and 7,060 donors in an hour.

The computational demand of scIntegral grows linearly respect to the size of the data. PBMC 68k(16) is one of the largest human scRNA-seq dataset available consisting 68,579 cells from human peripheral blood. We down-sampled the dataset and applied scIntegral to 5k, 10k, 20k, 40k and 68k cells. scIntegral on a single GPU took only 5.5, 6.9, 10.8 and 16.7 seconds respectively (**Figure 3d**). On a 4-core CPU, it took 67, 133, 327 and 678 seconds.

When the size of the data increases, the number of cells is not the only source of increased computational burden. Increased number of donors demands additional covariates to be incorporated into the model. To measure the impact of covariates, we augmented artificial covariates information to the PBMC 68k dataset. Even when the number of covariates increased, the runtime remained relatively constant in our method (**Supplemental_Table_S5.xlsx**). This showed that because of our efficient gradient implementation, scIntegral can handle multiple covariates seamlessly.

### scIntegral outperforms existing semi-supervised methods in classical tasks

We compare the classification accuracy of scIntegral to existing semi-supervised methods: SCINA and CellAssign. Human embryogenic stem cell (hESC) dataset and PBMC 4k dataset are both FACS sorted datasets with supposedly true cell type labels. When applied to these datasets, scIntegral showed superior accuracy compared to other two methods (**Figure 4a and 4b**). In the hESC data of Koh et al., scIntegral showed 96.8% median accuracy. In 20 different runs, this ranged from 96.4% to 96.9%. Thus, scIntegral showed robust performance not depending on random start. By contrast, SCINA and CellAssign achieved only 80.0% and 89.7% median accuracy, respectively. Across 20 runs, SCINA’s accuracy was constant, because the method is deterministic. However, CellAssign’s accuracy largely varied from 64.4% to 94.4% (**Figure 4a**). In the PBMC 4k data of Zheng et al., scIntegral achieved 97.0% median accuracy. Across 20 different runs, scIntegral gave an accuracy ranging from 96.9% to 97.0%. SCINA and CellAssign achieved 94% and 74.6% median accuracy, respectively. Across 20 runs, SCINA’s accuracy was constant and CellAssign’s accuracy largely varied from 25.0% to 95.9% (**Figure 4b**).

**Figure 4.**
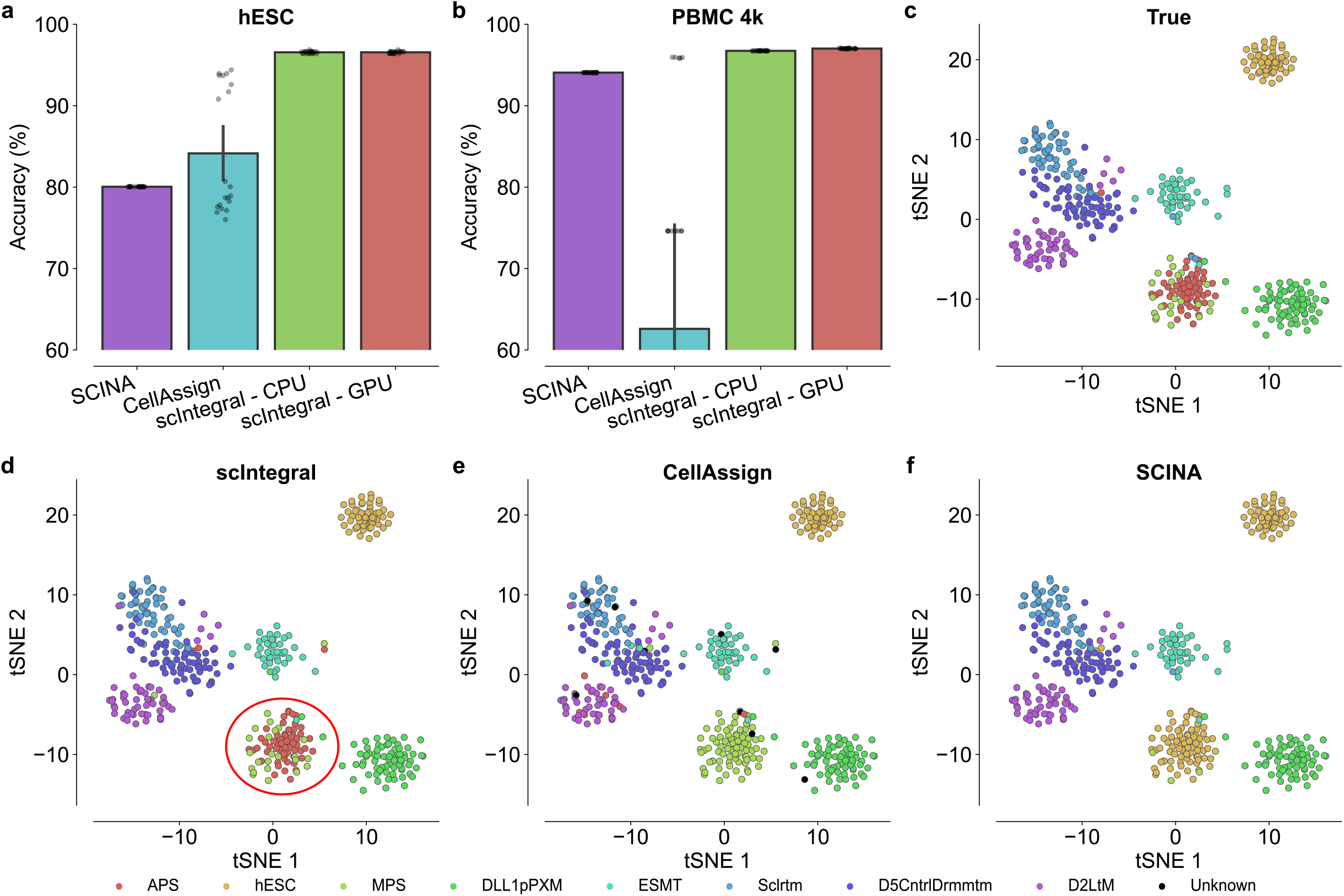
Accuracy benchmark of scIntegral and competing methods. **a.** Accuracy of SCINA(1), CellAssign and scIntegral in the hESC dataset(3). **b.** Accuracy of SCINA, CellAssign and scIntegral in the PBMC 4k dataset(16). **c.** t-SNE plot of the hESC dataset labeled by the true cell type labels. **d.** t-SNE plot of the hESC dataset labeled by assignment result of scIntegral. **e.** t-SNE plot of the hESC dataset labeled by assignment result of CellAssign. **f.** t-SNE plot of the hESC dataset labeled by assignment result of SCINA.

High precision of scIntegral is highlighted in the hESC dataset. Anterior primitive streak (APS) and mid primitive streak (MPS) are closely related cells that emerged during human developmental process(3). scIntegral successfully distinguished the two populations but SCINA and CellAssign did not (**Figure 4b–4d**). Looking at the cells in the red circles in the t-SNE plot, the MPS population colored in green are surrounding the APS population colored in red (**Figure 4b**). Since one population is distributed around the other in a circular manner, manual curation through vision must have been impossible. This shows that scIntegral can help identify similar populations even when they are distributed closely in the dimension reduced 2D plane.

## Discussion

We presented scIntegral, a cell-type classifier method that can integrate data of thousands of donors efficiently. Our method is a semi-supervised method that takes raw expression expression counts and marker signatures as inputs. scIntegral showed high accuracy in real data analysis and showed surprisingly high precision in identifying very rare cell populations. Importantly, we showed that integrating data from multiple donors can increase cell-type identification accuracy. scIntegral properly handles data heterogeneity which may confound the classification results.

scIntegral is flexible in the sense that it only requires marker signatures. Our semi-supervised classifier overcomes major limitations of existing supervised classifiers that require reference data. A drawback of supervised approaches is that they can only provide classification results based on a fixed set of cell-type labels since it is trained on a pre-labeled reference data. Additionally, reference data may not be available for the cell types in the sample. Another drawback is that there can be biological and technical heterogeneity between the reference data and the sample, which may affect the accuracy. scIntegral is free from these caveats because it is a reference-free semi-supervised method.

scIntegral systematically accounts for confounding by fully integrating covariates in the likelihood model. Other possible approaches to correct for confounding include normalization. To remove unwanted technical variations among different datasets, many existing approaches such as supervised approaches depend on normalization applied to each dataset. However, studies show that such normalization can remove meaningful biological signals which can degrade the classification quality(8,18). scIntegral overcomes this problem by directly modeling the raw expression count through a linear model that can incorporate multiple technical covariates.

Another famous cell-type classification regime is cluster-based labeling(14). In this regime, cells are primarily grouped based on clustering algorithms and then labeled altogether as a whole based on its mean characteristics. This procedure ignores the individual characters of the cell and exhibits low resolution(22). In contrast, scIntegral fully leverages the individual expression profile of each cell and uses them to provide accurate cell-type labels.

scIntegral also showed superior performance in classifying small conventional datasets. Especially, in the hESC dataset, scIntegral successfully distinguished developmentally close cell populations that were impossible to discern through visual information nor other existing semisupervised methods. Therefore, scIntegral has a high utility not only in large datasets but also in relatively small datasets that are routinely analyzed nowadays.

The size of scRNA-seq data are growing rapidly and now starts to reach consortium-level scale(22). Through scIntegral, we have demonstrated the benefits of aggregating multiple data by explicitly showing that classification accuracy is improved by integrating multiple donors. We expect that scIntegral will play an important role in an era of such large-scale data for the following reasons. First, it is highly scalable and can integrate thousands of donors in less than an hour. Second, it can account technical variations in a flexible manner. Third, it can provide accurate cell-type labels and performs well even when the candidate cell population is extremely rare.

## Data availability

scIntegral is an open source software and can be found at https://github.com/hanbin973/scIntegral.

## Acknowledgements

Not applicable

## Authors’ contributions

HL, CK, and BH devised the project. HL and CK processed the datasets and performed analyses. HL and CK wrote the software code. HL derived the gradient formulae. BH supervised the project. HL, CK, JJ, KJ, and BH wrote the manuscript. All authors read and approved the final manuscript.

## Reference

1. Zhang, Z., Luo, D., Zhong, X., Choi, J.H., Ma, Y., Wang, S., Mahrt, E., Guo, W., Stawiski, E.W., Modrusan, Z. et at. (2019) SCINA: A Semi-Supervised Subtyping Algorithm of Single Cells and Bulk Samples. Genes (Basel), 10.

2. Zhang, A.W., O’Flanagan, C., Chavez, E.A., Lim, J.L.P., Ceglia, N., McPherson, A., Wiens, M., Walters, P., Chan, T., Hewitson, B. et al. (2019) Probabilistic cell-type assignment of single-cell RNA-seq for tumor microenvironment profiling. Nat Methods, 16, 1007–1015.

3. Koh, P.W., Sinha, R., Barkal, A.A., Morganti, R.M., Chen, A., Weissman, I.L., Ang, L.T., Kundaje, A. and Loh, K.M. (2016) An atlas of transcriptional, chromatin accessibility, and surface marker changes in human mesoderm development. Sci Data, 3, 160109.

4. Muraro, M.J., Dharmadhikari, G., Grun, D., Groen, N., Dielen, T., Jansen, E., van Gurp, L., Engelse, M.A., Carlotti, F., de Koning, E.J. et al. (2016) A Single-Cell Transcriptome Atlas of the Human Pancreas. Cell Syst, 3, 385–394 e383.

5. Segerstolpe, A., Palasantza, A., Eliasson, P., Andersson, E.M., Andreasson, A.C., Sun, X., Picelli, S., Sabirsh, A., Clausen, M., Bjursell, M.K. et al. (2016) Single-Cell Transcriptome Profiling of Human Pancreatic Islets in Health and Type 2 Diabetes. Cell Metab, 24, 593–607.

6. Stuart, T., Butler, A., Hoffman, P., Hafemeister, C., Papalexi, E., Mauck, W.M., 3rd, Hao, Y, Stoeckius, M., Smibert, P. and Satija, R. (2019) Comprehensive Integration of Single-Cell Data. Cell, 177, 1888–1902 e1821.

7. Johansen, N. and Quon, G. (2019) scAlign: a tool for alignment, integration, and rare cell identification from scRNA-seq data. Genome Biol, 20, 166.

8. Korsunsky, I., Millard, N., Fan, J., Slowikowski, K., Zhang, F., Wei, K., Baglaenko, Y., Brenner, M., Loh, P.R. and Raychaudhuri, S. (2019) Fast, sensitive and accurate integration of singlecell data with Harmony. Nat Methods, 16, 1289–1296.

9. Bacher, R., Chu, L.F., Leng, N., Gasch, A.P., Thomson, J.A., Stewart, R.M., Newton, M. and Kendziorski, C. (2017) SCnorm: robust normalization of single-cell RNA-seq data. Nat Methods, 14, 584–586.

10. Hafemeister, C. and Satija, R. (2019) Normalization and variance stabilization of single-cell RNA-seq data using regularized negative binomial regression. Genome Biol, 20, 296.

11. Amrhein, L., Harsha, K. and Fuchs, C. (2019) A mechanistic model for the negative binomial distribution of single-cell mRNA counts. bioRxiv.

12. Townes, F.W., Hicks, S.C., Aryee, M.J. and Irizarry, R.A. (2019) Feature selection and dimension reduction for single-cell RNA-Seq based on a multinomial model. Genome Biol, 20, 295.

13. Lun, A.T., McCarthy, D.J. and Marioni, J.C. (2016) A step-by-step workflow for low-level analysis of single-cell RNA-seq data with Bioconductor. F1000Res, 5, 2122.

14. Butler, A., Hoffman, P., Smibert, P., Papalexi, E. and Satija, R. (2018) Integrating single-cell transcriptomic data across different conditions, technologies, and species. Nat Biotechnol, 36, 411–420.

15. Franzen, O., Gan, L.M. and Bjorkegren, J.L.M. (2019) PanglaoDB: a web server for exploration of mouse and human single-cell RNA sequencing data. Database (Oxford), 2019.

16. Zheng, G.X., Terry, J.M., Belgrader, P., Ryvkin, P., Bent, Z.W., Wilson, R., Ziraldo, S.B., Wheeler, T.D., McDermott, G.P., Zhu, J. et at. (2017) Massively parallel digital transcriptional profiling of single cells. Nat Commun, 8, 14049.

17. Svensson, V. (2020) Droplet scRNA-seq is not zero-inflated. Nat Biotechnol, 38, 147–150.

18. Kim, T.H., Zhou, X. and Chen, M. (2020) Demystifying “drop-outs” in single-cell UMI data. Genome Biol, 21, 196.

19. Love, M.I., Huber, W. and Anders, S. (2014) Moderated estimation of fold change and dispersion for RNA-seq data with DESeq2. Genome Biol, 15, 550.

20. Robinson, M.D., McCarthy, D.J. and Smyth, G.K. (2010) edgeR: a Bioconductor package for differential expression analysis of digital gene expression data. Bioinformatics, 26, 139–140.

21. MacParland, S.A., Liu, J.C., Ma, X.Z., Innes, B.T., Bartczak, A.M., Gage, B.K., Manuel, J., Khuu, N., Echeverri, J., Linares, I. et al. (2018) Single cell RNA sequencing of human liver reveals distinct intrahepatic macrophage populations. Nat Commun, 9, 4383.

22. van der Wijst, M., deVries, D.H., Groot, H.E., Trynka, G., Hon, C.C., Bonder, M.J., Stegle, O., Nawijn, M.C., Idaghdour, Y., van der Harst, P. et al (2020) The single-cell eQTLGen consortium. Elife, 9.

