## Supplementary Note for "scIntegral: A scalable and accurate cell-type identification method for scRNA-seq data with application to integration of multiple donors"

### scIntegral: Supplementary Note

#### 1 Negative binomial and quasi-Poisson regression

Negative binomial regression and quasi-Poisson regression are two famous regression methods for overdispersed count data. In our study we used the `glm.nb` from `MASS` package and `glm` from R base. The regression model fitted is as follows.

$$\log \mathbb{E} y_g = \beta_0 + \sum_p \beta_{gp} x_{np} + \sum_t \gamma_{gt} \cdot \mathbb{I}_t + s$$

Here, $\beta_0$ is the constant, $\beta_{gp}$ are covariate coefficients, $\gamma_{gt}$ are cell-type specific expression coefficient and $s$ is the size-factor. $x$ and $\mathbb{I}$ are covariate and cell-type indicators respectively.

#### 2 Derivation of Gradient

Here we describe the detailed derivation of the gradient of our likelihood function. As described in **Online Methods**, we model the mean UMI count $\mu_{ngt}$ of gene $g$ in the cell $n$ of cell-type $t$ as

$$\log \mu_{ngt} = \beta_{g0} + \Sigma_p \beta_{gp} x_{np} + \delta_{gt} m_{gt} + \log s_n$$

where $\beta_{g0}$ is the baseline expression of gene $g$, $\beta_{gp}$ is the effect of covariate $p$ on gene $g$, $x_{np}$ is the value of covariate $p$ in cell $n$, $\delta_{gt}$ is the gene expression change specific to cell-type $t$, $m_{gt}$ is a binary variable indicating whether $g$ is a marker for cell-type $t$, and $s_n$ is the pre-calculated cell size factor.

Let $y_{ng}$ be the observed UMI count of gene $g$ in cell $n$. We assume that $y_{ng}$ follows a negative-binomial distribution.

$$y_{ng} \sim NB(\mu_{ngt}, \phi_g)$$

Existing theoretical literature predict that the dispersion of the expression count is determined by the gene regulation mechanism. By assuming that gene regulation mechanism is independent of the celltype for the same gene, we model the dispersion to be dependent only on the gene $g$ but not celltype $t$, thus $\phi_g$.

The full likelihood is

$$LL = \sum_n \log\left[\sum_t \exp\left(\sum_g \log NB(y_{ng}; \mu_{ngt}, \phi_g)\right)\right]$$

Before proceeding, we calculate the first derivative of the negative-binomial log probability density function.

$$\begin{aligned}\log NB(y;\mu,\phi) &= \log\Gamma(y+\phi) - \log\Gamma(y+1) - \log\Gamma(\phi)\\ &+ y\left(\log(\mu) - \log(\mu+\phi)\right)\\ &+ \phi\left(\log(\phi) - \log(\mu+\phi)\right) \qquad (1)\end{aligned}$$

Therefore,

$$\begin{aligned}\frac{\partial}{\partial\mu}\log NB(y;\mu,\phi) &= y\cdot\frac{1}{\mu} - y\cdot\frac{1}{\mu+\phi} - \phi\cdot\frac{1}{\mu+\phi} \qquad (2)\\ &= \frac{\phi(y-\mu)}{\mu(\mu+\phi)} \qquad (3)\end{aligned}$$

and

$$\begin{aligned}\frac{\partial}{\partial\phi}\log NB(y;\mu,\phi) &= \psi(y+\phi) - \psi(\phi) - \frac{y}{\mu+\phi}\\ &+ \log(\phi) + 1 - \log(\mu+\phi) - \frac{\phi}{\mu+\phi}\\ &= \psi(y+\phi) - \psi(\phi) + \log\left(\frac{\phi}{\phi+\mu}\right) + \frac{\mu-y}{\mu+\phi} \qquad (4)\end{aligned}$$

Now we calculate the derivative of $LL$

$$\frac{\partial LL}{\partial \delta_{gt}} = \sum_n \frac{\partial \left[\sum_t \exp\left(\sum_g \log NB(y_{ng};\mu_{ngt},\phi_g)\right)\right] / \partial \delta_{gt}}{\sum_{t'} \exp\left(\sum_g \log NB(y_{ng};\mu_{ngt'},\phi_g)\right)}$$

Cell types other than $t$ vanish in the numerator,

$$= \sum_n \frac{\partial \left[\exp\left(\sum_g \log NB(y_{ng};\mu_{ngt},\phi_g)\right)\right] / \partial \delta_{gt}}{\sum_{t'} \exp\left(\sum_g \log NB(y_{ng};\mu_{ngt'},\phi_g)\right)}$$

$$= \sum_n \frac{\left[\exp\left(\sum_g \log NB(y_{ng};\mu_{ngt},\phi_g)\right)\right] \partial \left(\sum_g \log NB(y_{ng};\mu_{ngt},\phi_g)\right) / \partial \delta_{gt}}{\sum_{t'} \exp\left(\sum_g \log NB(y_{ng};\mu_{ngt'},\phi_g)\right)}$$

Genes other than $g$ can be considered constant,

$$= \sum_n \frac{\exp\left(\sum_g \log NB(y_{ng};\mu_{ngt},\phi_g)\right)}{\sum_{t'} \exp\left(\sum_g \log NB(y_{ng};\mu_{ngt'},\phi_g)\right)} \cdot \frac{\partial \left(\log NB(y_{ng};\mu_{ngt},\phi_g)\right)}{\partial \delta_{gt}}$$

by equation (3),

$$= \sum_n \frac{\exp\left(\sum_g \log NB(y_{ng};\mu_{ngt},\phi_g)\right)}{\sum_{t'} \exp\left(\sum_g \log NB(y_{ng};\mu_{ngt'},\phi_g)\right)} \cdot \frac{\phi_g(y_{ng}-\mu_{ngt})}{\mu_{ngt}(\mu_{ngt}+\phi_g)} \cdot \frac{\partial \mu_{ngt}}{\partial \delta_{gt}}$$

$$= \sum_n \frac{\exp\left(\sum_g \log NB(y_{ng};\mu_{ngt},\phi_g)\right)}{\sum_{t'} \exp\left(\sum_g \log NB(y_{ng};\mu_{ngt'},\phi_g)\right)} \cdot \frac{\phi_g(y_{ng}-\mu_{ngt})}{\mu_{ngt}(\mu_{ngt}+\phi_g)} \cdot \mu_{ngt} \cdot 1$$

$$= \sum_n \frac{\exp\left(\sum_g \log NB(y_{ng};\mu_{ngt},\phi_g)\right)}{\sum_{t'} \exp\left(\sum_g \log NB(y_{ng};\mu_{ngt'},\phi_g)\right)} \cdot \frac{\phi_g}{\mu_{ngt}+\phi_g} \cdot (y_{ng}-\mu_{ngt})$$

$$\frac{\partial LL}{\partial \phi_g} = \sum_n \frac{\partial \left[\sum_t \exp\left(\sum_g \log NB(y_{ng}; \mu_{ngt}, \phi_g)\right)\right] / \partial \phi_g}{\sum_{t'} \exp\left(\sum_g \log NB(y_{ng}; \mu_{ngt'}, \phi_g)\right)}$$

$$= \sum_n \frac{\sum_t \left[\partial \exp\left(\sum_g \log NB(y_{ng}; \mu_{ngt}, \phi_g)\right) / \partial \phi_g\right]}{\sum_{t'} \exp\left(\sum_g \log NB(y_{ng}; \mu_{ngt'}, \phi_g)\right)}$$

$$= \sum_n \sum_t \frac{\left[\exp\left(\sum_g \log NB(y_{ng}; \mu_{ngt}, \phi_g)\right)\right] \cdot \partial \left[\sum_g \log NB(y_{ng}; \mu_{ngt}, \phi_g)\right] / \partial \phi_g}{\sum_{t'} \exp\left(\sum_g \log NB(y_{ng}; \mu_{ngt'}, \phi_g)\right)}$$

Genes other than $g$ can be considered constant,

$$= \sum_n \sum_t \frac{\left[\exp\left(\sum_g \log NB(y_{ng}; \mu_{ngt}, \phi_g)\right)\right] \cdot \partial \left[\log NB(y_{ng}; \mu_{ngt}, \phi_g)\right] / \partial \phi_g}{\sum_{t'} \exp\left(\sum_g \log NB(y_{ng}; \mu_{ngt'}, \phi_g)\right)}$$

by equation (4),

$$= \sum_n \sum_t \frac{\exp\left(\sum_g \log NB(y_{ng}; \mu_{ngt}, \phi_g)\right)}{\sum_{t'} \exp\left(\sum_g \log NB(y_{ng}; \mu_{ngt'}, \phi_g)\right)}$$

$$\times \left[\psi(y_{ng} + \phi_g) - \psi(\phi_g) + \log\left(\frac{\phi_g}{\phi_g + \mu_{ngt}}\right) + \frac{\mu_{ngt} - y_{ng}}{\mu_{ngt} + \phi_g}\right]$$

$$\frac{\partial LL}{\partial \beta_{gp}} = \sum_n \frac{\partial \left[\sum_t \exp\left(\sum_g \log NB(y_{ng}; \mu_{ngt}, \phi_g)\right)\right] / \partial \beta_{gp}}{\sum_{t'} \exp\left(\sum_g \log NB(y_{ng}; \mu_{ngt'}, \phi_g)\right)}$$

$$= \sum_n \sum_t \frac{\exp\left(\sum_g \log NB(y_{ng}; \mu_{ngt}, \phi_g)\right)}{\sum_{t'} \exp\left(\sum_g \log NB(y_{ng}; \mu_{ngt'}, \phi_g)\right)} \cdot \frac{\phi_g}{\mu_{ngt} + \phi_g} \cdot (y_{ng} - \mu_{ngt}) \cdot x_{np}$$

Thus, we have successfully derived the derivative form in **Online Methods**.
